# Temporally variable drug profiles select for diverse adaptive pathways despite conservation of efflux-based resistance mechanism

**DOI:** 10.1101/2023.05.19.541537

**Authors:** Akanksha, Sarika Mehra

## Abstract

Antibiotic resistance is a global health concern with emergence of resistance in bacteria out-competing the discovery of novel drug candidates. While Adaptive Laboratory Evolution (ALE) has been used to identify bacterial resistance determinants, most studies investigate evolution under stepwise increasing drug profiles. Thus, bacterial adaptation under long-term constant drug concentration, a physiologically relevant profile, remains underestimated. Using ALE of *Mycobacterium smegmatis* subjected to a range of Norfloxacin concentrations under both constant and stepwise increasing drug dosage, we investigated the impact of variation of drug profiles on resistance evolution. All the evolved mutants exhibited a drug concentration dependent increase in resistance accompanied with an increase in the number of mutations. Mutations in an efflux pump regulator, LfrR, were found in all the evolved populations suggesting conservation of an efflux-based resistance mechanism. The selection of these mutations was tightly coupled to the presence of its regulated gene in the genetic background. Further, *lfr*R mutations appeared early during the adaptive trajectory and imparted low-level resistance. Subsequently, sequential acquisition of other mutations, dependent on the drug profile, led to high-level resistance emergence. While divergent mutational trajectories led to comparable phenotype, populations evolved under constant drug exposure accumulated mutations in dehydrogenase genes whereas in populations under increasing drug exposure, mutations in additional regulatory genes were selected. Our data also shows that irrespective of the evolutionary trajectory, drug target mutations were not selected up to 4X drug concentration. Overall, this work demonstrates that evolutionary trajectory is strongly influenced by the drug profile.

## Importance

Since their discovery, antibiotics have proved to be a boon for mankind. However, due to the uncontrolled use of antibiotics, bacteria are rapidly developing antibiotic resistance rendering them ineffective. The rate of antibiotic resistance emergence in bacteria is very fast compared to the rate of discovery of new antibiotics compelling us to find newer ways to predict and control antibiotic resistance. While certain resistance mechanisms can be overcome through the use of adjuvants along with the drug, others render the antibiotic ineffective. In this work, we use ALE to determine how the variation in drug profile that the bacteria are exposed to, impacts the mechanism of resistance selected. We show that an efflux-based resistance mechanism is conserved in all the evolved populations and appears early in the evolutionary trajectory. Subsequently, the evolutionary trajectory diverges depending on the drug profile. While populations evolved under a constant drug regime accumulate mutations in metabolic genes, those under a stepwise increasing drug concentration, harbor mutations in additional regulatory genes. The data also shows that irrespective of the drug profile, mutations in drug target genes are absent in all evolved populations up to 4X norfloxacin MIC concentration. Knowledge of adaptive pathway under diverse drug profiles would help to design effective treatment strategies.

## Introduction

Antibiotics remain one of the most significant discoveries in the field of medicine. However, the optimism associated with antibiotic discovery has started to decline owing to the rapid emergence of antibiotic resistance in almost all human associated pathogens. The emergence of Antimicrobial Resistance (AMR) also leads to a global economic burden on healthcare facilities (1). Antibiotic treatment can have two conflicting effectsdesired immediate inhibition of bacterial growth and an undesired effect of promoting resistance evolution (2). Unfortunately, the indiscriminate use of these wonder medicines has led to rapid emergence of resistant strains (3). According to the U.K task force, by 2050, 10 million deaths are estimated worldwide due to antibiotic resistant infections (4).

In order to predict and control bacterial resistance emergence, identification and understanding of constraints shaping the evolution of resistance is critical (5). There are multiple mechanisms through which bacteria attain resistance, such as mutations in the drug target genes leading to reduced binding of the drug, altered drug uptake and efflux through expression of efflux pumps (6, 7). Understanding which of these mechanisms will evolve under a given drug regime, can guide the development of targeting strategies. Adaptive laboratory evolution (ALE) proves to be a very useful tool for gaining these insights and thus has been employed for different pathogens in the past (8).

The evolution of resistance is governed by many intrinsic attributes such as the genetic background of the organism (9, 10), spontaneous mutation rates and the fitness effects of these mutations both in the presence as well as in the absence of the selection pressure (11, 12). Apart from the contribution of bacterial genetics, ecology also plays a crucial role. The antibiotic selection pressure is the major contributor of resistance acquisition and directly selects for mutations that impart resistance. Under inhibitory concentrations of the drug, any mutation providing resistance and survival is selected with fitness being secondary whereas under non-inhibitory concentrations, fitness becomes critical and mutations with greater fitness advantage are selected (13, 14). Many other selection parameters further influence the evolutionary outcomes. Evolution under randomly fluctuating environments has been shown to lead to evolved populations with increased fitness against novel stresses due to increased efflux activity in the selected populations (15). Recent studies have also explored the role of metabolism on the adaptive outcome. For example, *E. coli* evolved under two different carbon sources (glucose vs acetate), led to rapid resistance in cells cultured on glucose, due to the increased expression of efflux pumps allowing switch from respiration to fermentation which was absent when cells were grown in acetate (16). Another important factor is the mutational supply. Drug-target mutations usually have a higher fitness cost and are rare and thus are selected only in case of high transfer size. In contrast, evolution at lower population bottlenecks yields many mutations with low fitness (17). Thus, apart from affecting whether resistance will emerge, the combination of intrinsic and environmental parameters also influence which mechanisms will appear. Overall, bacterial resistance evolution is a complex phenomenon involving multiple players.

Fluoroquinolones (FQs) are broad-spectrum antibiotics used for the treatment of multiple bacterial infections. These drugs primarily target DNA gyrase and DNA topoisomerase IV leading to defective replication and ultimately death (18). Drug target mutations and efflux pump over-expression are the two most common resistance mechanisms for fluoroquinolone drugs (19–21). While mutations in drug target genes are responsible for most of the resistance in clinical strains, about 20-30% of clinical strains have been reported to not contain any of the known target gene mutations (22, 23). Recent studies have reported efflux pumps to be over-expressed in many clinical strains (21, 24, 25). While in some strains the efflux pumps are constitutively over-expressed, in other strains they are induced upon drug treatment (26, 27), suggesting mutations in the associated transcriptional regulators.

FQ resistance can be easily selected in laboratory settings by serial passaging with varying resistance levels of the selected mutants (28–30). However, most ALE studies are carried out under conditions that allow for the selection of maximum resistance, generally via drug target mutations, such as under stepwise increasing drug concentration or higher transfer size (10, 17, 28, 31–33). The effect of other adaptive regimes, such as long-term exposure to a drug concentration, is not well studied. The pharmacokinetics of drugs in the host is likely to experience a constant drug pressure with intermediate periods of low drug concentration (34, 35). The stepwise increasing drug profiles are less physiologically relevant.

In this study, we systematically investigate the effect of drug profiles on the progression of resistance evolution using *M. smegmatis* as a model system and Norfloxacin as a representative drug. We restrict the adaptive regimes to single colony and medium transfer sizes that may be physiologically more relevant (36, 37). We study how temporally constant drug profiles, over several generations, ranging from sub-MIC to 4X MIC concentrations affect the evolutionary trajectories. We also compare these to the stepwise increasing drug profiles both in terms of the genotype and the phenotype of the evolved populations. Knowledge of these factors would be critical in devising effective strategies under different situations to counter antibiotic resistance.

## Results

### Emergence of mutants with resistance levels proportional to the selection strength

We first consider the effect of a constant drug pressure, ranging from sub-MIC to 4X MIC, on the evolution of AMR. ALE was carried out for wild-type *Mycobacterium smegmatis* under different constant drug profiles where the given drug concentration was held constant for more than 300 generations. The Norfloxacin concentration ranged from 0.5X MIC to 4X MIC (in duplicates) giving rise to eight independent drug evolved lineages. Parallelly, the WT was also passaged in the absence of the drug for a similar number of generations. At the end of the evolution experiment, six independent end point isolates (3 from each replicate) were selected for each drug evolved lineage. Evolution profile for the different Constant populations has been represented in Figure 1.A.

**Figure 1.**
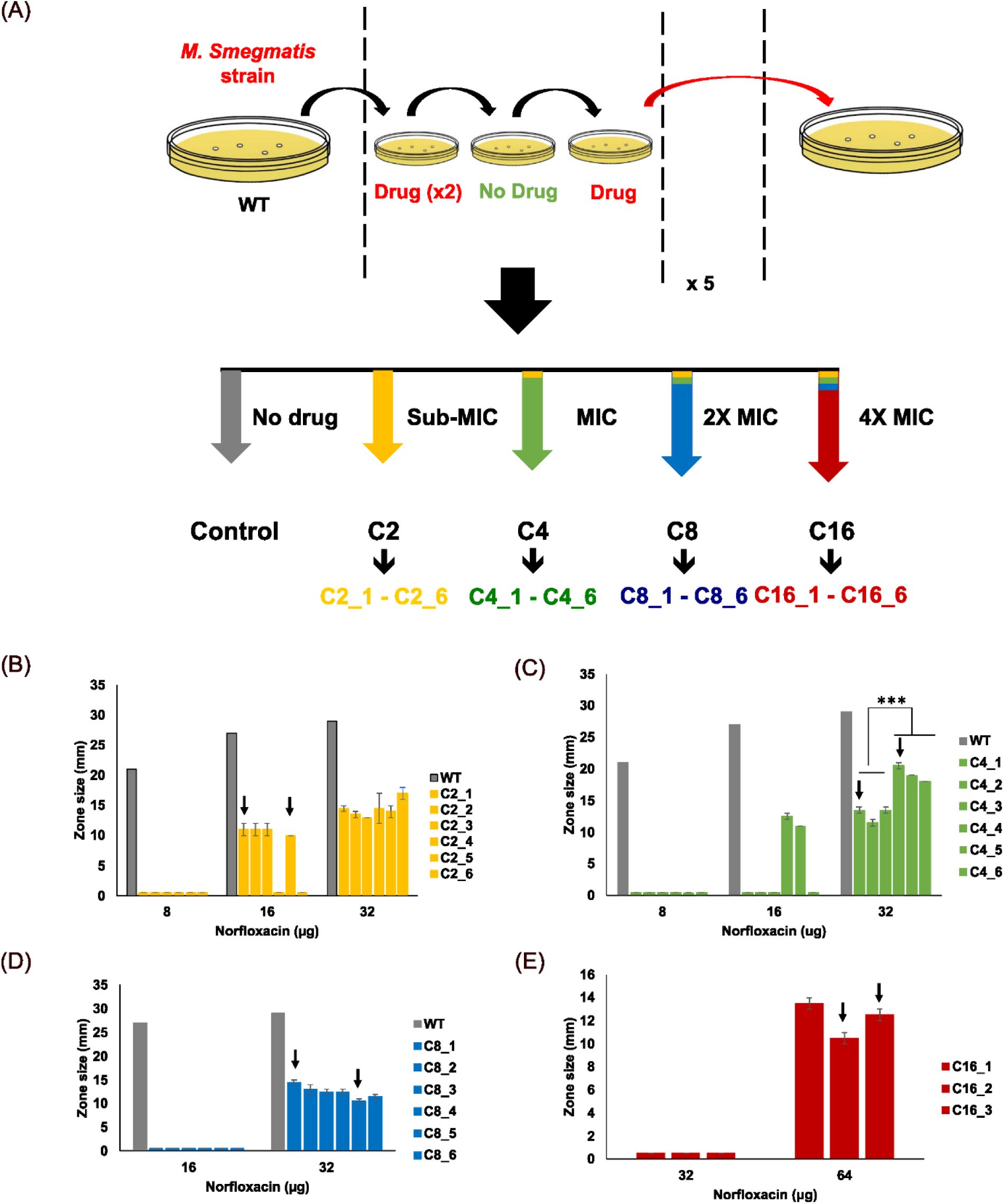
(A) Evolution profile of the different Constant populations. The first letter C represents the constant drug profile followed by the drug concentration (2-16 µg/ml) during evolution. The last digit denotes the colony number. (B), (C), (D) and (E) show Norfloxacin sensitivity of Constant 2 (C2), Constant 4 (C4), Constant 8 (C8) and Constant 16 (C16) populations respectively. The Y-axis represents the zone size in millimeters (mm) and the X-axis represents the Norfloxacin amount (μg). Standard deviation has been calculated over three replicates. * Indicates P < 0.05, ** indicates P < 0.01 and *** indicates P < 0.001. The colonies selected for further characterization have been indicated by an arrow.

The six end point colonies from each lineage were further characterized with respect to their susceptibility to Norfloxacin, the selection pressure (details in methods). For C16 populations, data is available for only three colonies out of the six colonies selected as the rest did not grow. The evolved populations exhibited varying levels of resistance to Norfloxacin ranging from 4X to 16X increase (Figure 1.B to 1.E). All the individual colonies from a population exhibited similar sensitivity except the C4 population where the two replicate lineages displayed significant differences in sensitivity. A maximum of two representative colonies from each of the evolved populations were selected for further characterization such that the diversity among the populations is captured well.

### Number of mutations selected is proportional to the selection strength

Evolution under constant drug selection pressure yielded mutants with distinct resistance levels against the selection pressure. The increase in resistance was proportional to the strength of the selection pressure. The C2 population exhibited a 4-fold increase in MIC, followed by an 8-fold increase for the C4 and C8 populations. The C16 population exhibited the highest MIC with a 16-fold increase in Norfloxacin MIC. Complete killing of the C16 population was not achieved even at 32X MIC (Figure 2.A and 2.B). The increased resistance however had a fitness cost and led to an increase in doubling time of the evolved populations. As seen in Figure 2.C, among different populations, doubling time ranged from 2.9 hours to 3.2 hours with respect to WT doubling time of 2.5 hours. Significant increase in lag time of greater than 5 hours was observed in C8 and C16 populations with respect to the WT strain (Figure 2.D).

**Figure 2.**
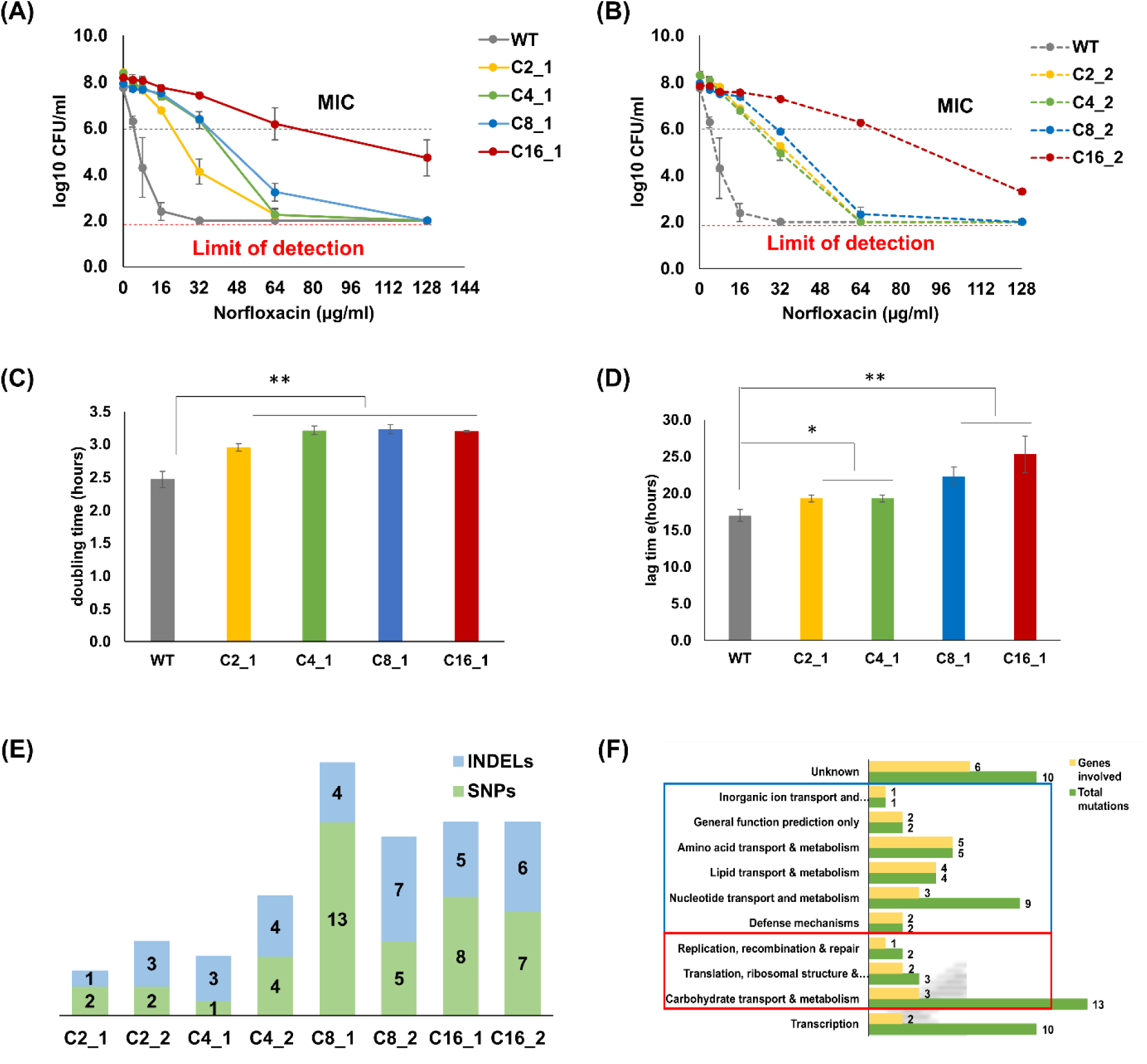
Phenotypic and genotypic characterization of the two representative constant colonies. (A) and (B) Change in cell viability at different Norfloxacin concentrations. Viability was determined using the broth microdilution method followed by CFU count (Limit of detection: 10^2^ CFU/ml, initial cell count: 10^6^ CFU/ml). Y-axis represents cell density while the X-axis represents Norfloxacin concentration. (C) and (D) represent doubling time and lag time respectively for the evolved constant colonies. Error bars have been calculated over triplicate samples. * Indicates P < 0.05, ** indicates P < 0.01 and *** indicates P < 0.001. (E) Number of unique mutations identified in different colonies of the evolved populations with respect to the Control population. Filter of frequency>0.3 and DP>50 has been applied. (F) Distribution of mutations in the evolved populations according to functional Clusters of Orthologous Genes (COGs). Categories in the blue box are present in the C8 population while the ones in the red box are present in the C16 population.

To unravel the resistance determining factors, Whole Genome Sequencing (WGS) was carried out for each of the two representative colonies from the population (details in methods). Unique mutations in the evolved colonies were spread across the genome spanning both coding and non-coding regions. Overall, several unique mutations were identified in genes belonging to functional categories spanning DNA repair and replication, protein synthesis, and transport of nucleotides, lipids and amino acids to name a few. Surprisingly, there were no mutations in the drug target genes, *gyrA* and *gyrB* genes that encode for the gyrase protein subunits. Instead, mutation in a regulatory gene, MSMEG_6223 (*lfr*R) was conserved in all the evolved populations. Mutations in a hypothetical protein (MSMEG_1959) were also found to be conserved in multiple populations.

Interestingly, the number of mutations selected was proportional to the selection strength. While the C2 population had a maximum of 5 mutations, there were 13 mutations in the C16 population (Figure 2.E). Further, C8 and C16 populations had mutations in a diverse set of genes as compared to the C2 and C4 population (Figure 2.F). While the C8 population harbored mutations in genes involved in pathways related to transport and metabolism of nucleotides, amino acids and lipids, the C16 populations harbored mutations in genes involved in replication, recombination, and repair, translation, ribosomal structure and biogenesis and carbohydrate transport and metabolism. The complete list of all the unique mutations in the different populations with respect to Control population has been provided in Supplementary File S1. The distribution of the unique mutations in the different evolved colonies was quite diverse with very little overlap suggesting multiple adaptive trajectories.

### Contrasting drug profiles can lead to similar phenotype despite different mutational trajectory

Among different Constant populations, resistance increased as a function of selection strength. Also, the number of mutations selected increased for populations exhibiting higher resistance. Next, we wanted to compare the evolutionary trajectory of populations evolved under contrasting drug profiles but experiencing the same final selection strength. For this, we evolved WT *M. smegmatis* strain under stepwise increasing Norfloxacin concentration from sub-MIC to 4X WT MIC, referred to as the Ramp (R) population (Figure 3.A). From the end passage, three colonies, (R1, R2 and R3) were selected for further characterization and compared with C16_1 colony as a representative of the C16 population. The three Ramp colonies exhibited similar sensitivity to Norfloxacin as the C16_1 strain (Figure 3.B). Also, survival was similar for all the three Ramp colonies and C16_1 strain till Norfloxacin 64 µg/ml concentration. However, R1 strain displayed enhanced killing at Norfloxacin 128 µg/ml concentration (Figure 3.C). Thus, phenotypically Ramp population exhibited significant similarity to the C16 population despite experiencing contrasting drug profiles. The increased resistance in the Ramp colonies led to a decrease in fitness with respect to the WT strain in terms of both the doubling time and the lag time. However, when compared to the C16 population, no significant difference was observed (Supplementary file S2, Table S1).

**Figure 3.**
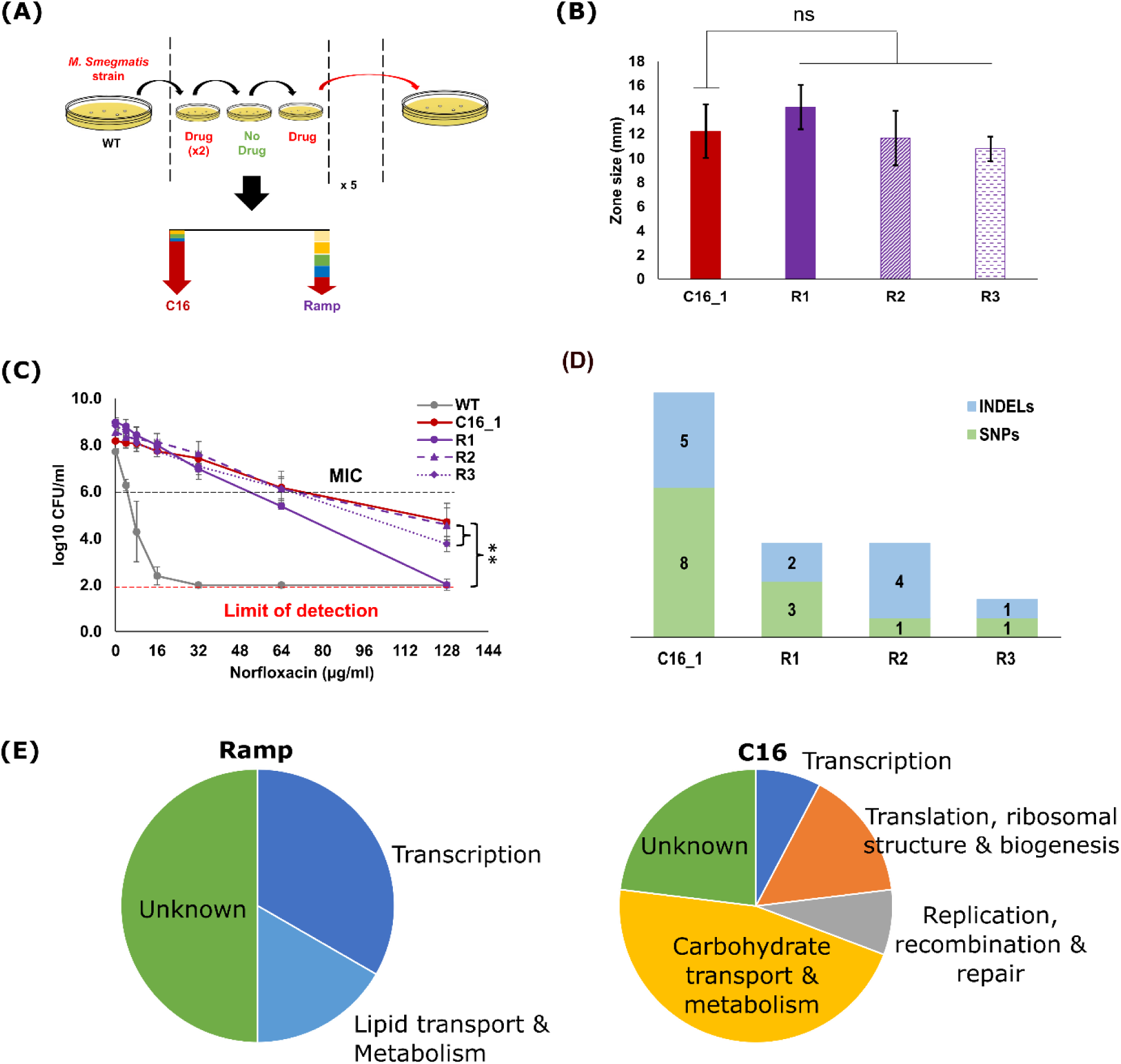
Phenotypic and genotypic characterization of the different evolved Ramp colonies. (A) Comparison of evolutionary profile for Ramp and C16 population. C16 evolution includes 3 passages under stepwise increase from Norfloxacin 2 μg/ml to Norfloxacin 8 μg/ml followed by 20 passages under constant Norfloxacin 16 μg/ml while Ramp evolution includes 20 passages under stepwise increasing concentration from Norfloxacin 1 μg/ml to Norfloxacin 16 μg/ml. (B) Comparison of susceptibility to Norfloxacin of selected colonies from the evolved Ramp and C16 population (Limit of detection: 10 mm). (C) Change in cell viability for the different Ramp colonies and C16_1 strain at different Norfloxacin concentrations as measured using broth dilution method followed by colony counting using Drop count method (Limit of detection: 10^2^ CFU/ml, initial cell count: 10^6^ CFU/ml). Error bars have been calculated using triplicate samples. * Indicates P < 0.05, ** indicates P < 0.01 and *** indicates P < 0.001. (D) Comparison of number of unique mutations identified in Ramp colonies and C16_1 strain. Filter of frequency > 0.3 and depth > 50 has been applied. (E) Distribution of mutations in Ramp and C16 population according to functional Clusters of Orthologous Genes (COGs).

WGS of the Ramp colonies revealed that the number of unique mutations in the Ramp population were significantly lower compared to the C16 population (Figure 3.D). Further, there was little overlap in the functional classes of the genes where mutations were present. In contrast to the C16 population, no mutations were seen in genes related to translation, replication, DNA repair or recombination in any of the Ramp colonies. On the contrary, many mutations were in and around regulatory genes (Figure 3.E). Similar to the previously evolved Constant populations, all the three Ramp colonies harbored a mutation in the regulatory gene, *lfr*R. In addition, a mutation in the intergenic region upstream of MSMEG_0573 (RpoE1) and MSMEG_0575 (Mmps1) was conserved in R1 and R3 colonies along with a mutation in MSMEG_5019 (regulatory protein) in R1. Drug target mutations were absent in these populations too. A list of all the unique mutations in Ramp population has been provided in Supplementary file S1.

### Diverse adaptive conditions affect the number of mutations but not the spectrum of mutations selected

Mutations in the *lfr*R gene were conserved in the populations that evolved under both constant and increasing drug pressure up-to 4X MIC of the drug, Norfloxacin. We next examine if alternate conditions such as growth in liquid media has any effect on the spectrum of mutations selected. Evolution was carried out in liquid media under a constant selection pressure of 2 µg/ml (sub-MIC) and 16 µg/ml (4X MIC) of Norfloxacin (Figure 4.A). These populations were referred to as C2_L and C16_L where C refers to the constant drug pressure, 2 and 16 represent the drug concentration used while L represents growth in liquid media during evolution. Evolution in liquid media also allowed for a higher transfer size of 10^5^ cells per passage and lower number of generations. Further, WT *M. smegmatis* strain was also evolved under stepwise increasing Norfloxacin concentration from sub-MIC to 4X MIC in liquid media and the evolved population was referred to as WT_Ev (Figure 4.A) with its evolution profile similar to the Ramp population.

**Figure 4.**
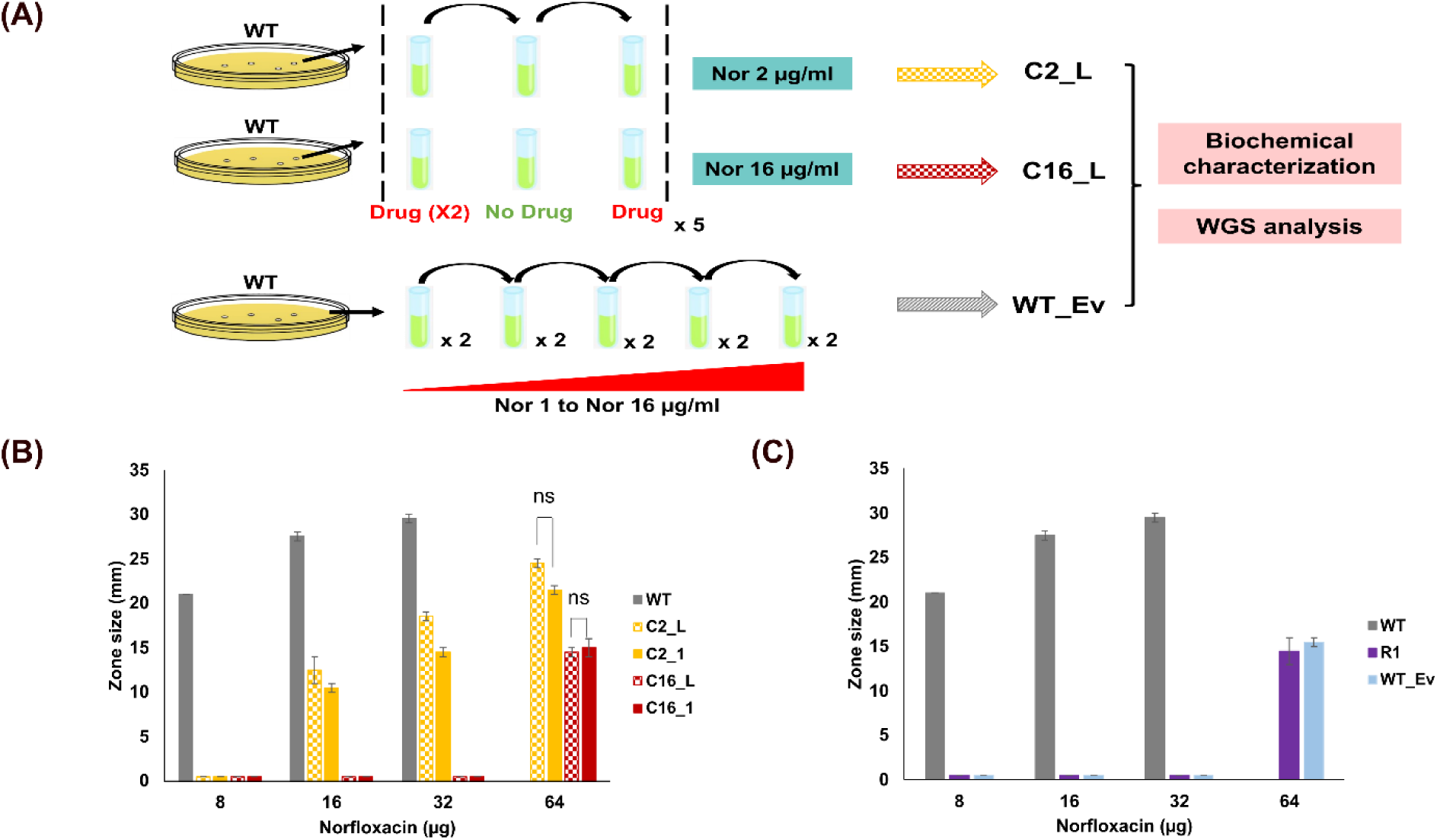
(A) Schematic for evolution of WT *M. smegmatis* under Constant and increasing Norfloxacin pressure in liquid growth medium. (B) and (C) Susceptibility of C2_L, C16_L and WT_Ev populations to Norfloxacin. The susceptibility of the C2_1, C16_1 and R1 colonies are shown for comparison respectively. Error bars have been calculated over triplicate samples.

Evolution in liquid media with higher transfer size allowed for direct growth in Norfloxacin 16 µg/ml which was not possible in solid media. However, the first passage on such high Norfloxacin concentration took around four days (96 hours). Sub-culturing was done every 20 hours for the C2_L population while every 22-24 hours for the C16_L population as opposed to 16 hours for the control population. For the WT_Ev population, sub-culturing was done every 24 hours except for the final passages at Norfloxacin 16 µg/ml requiring 48 hours.

Phenotypically, the liquid evolved populations resembled the solid evolved populations in the level of resistance acquired with a 4-fold increase for C2_L population and a 16-fold increase for C16_L and WT_Ev population (Figure 4.B and 4.C). However, the liquid evolved populations were more fit than the solid evolved populations and exhibited doubling time and lag time similar to the WT strain (Supplementary file S2, Table S2).

WGS of all the three evolved populations, C2_L, C16_L and WT_Ev revealed a lesser number of mutations when compared to the solid evolved populations. This lower number of mutations could partly be due to the lesser number of generations in the liquid evolved populations. Interestingly, even with a higher transfer size of 10^5^ cells, no mutations in drug target genes were seen. However, mutations in the *lfr*R gene were conserved. The C16_L population also harbored a deletion in the gene, MSMEG_5045 (D-2-hydroxyglutarate dehydrogenase) that was exactly the same as that in the C4_1 population. WT_Ev population harbored a mutation (SNP) in the LfrA efflux pump leading to change in Valine at the 378th position to Glycine (V378G) along with an insertion in the *lfr*R gene. A list of all the unique mutations in liquid evolved populations has been provided in Supplementary file S1.

### Conservation of a regulatory gene mutation causing over-expression of efflux pump despite different evolutionary trajectories of the evolved mutants

The various drug profiles yielded mutants with different resistance levels, fitness parameters and mutational spectrum (Table 1). These mutants were categorized into low and high resistance mutants with mutants exhibiting MIC of Norfloxacin higher than 4X as compared to the WT referred to as the high resistance mutants. Interestingly, most of the evolved mutants exhibited high resistance phenotype.

**Table 1.** Summary of phenotypic and genotypic attributes of the different evolved mutants. The high frequency mutations mentioned here are the ones above cut-off of frequency 0.8 and depth 50.

| Strain | Drug conc.<br>(µg/ml) | Nor MIC | EtBr MIC | Change in<br>Doubling time | Resistance level | High frequency mutations |  |  |
| --- | --- | --- | --- | --- | --- | --- | --- | --- |
|  |  |  |  |  |  | *lfrR* | Dehydrogenase | Others |
| WT | 0 | 4 | 7 | - | - | - | - | - |
| Constant drug profile |  |  |  |  |  |  |  |  |
| C2_1 | 2 | 16 | 14 | Increase | Low | Yes | No | MSMEG_0781,<br>MSMEG_4138 |
| C2_2 | 2 | 16 | 14 | Increase | Low | Yes | No | MSMEG_6459 |
| C2_L | 2 | 16 | 14 | No change | Low | Yes | No | None |
| C4_2 | 4 | 16 | 14 | Increase | Low | Yes | No | MSMEG_4902,<br>MSMEG_3427,<br>MSMEG_4383,<br>MSMEG_4514,<br>upstream of<br>MSMEG_1333 and<br>*hemE* |
| C4_1 | 4 | 32 | 28 | Increase | High | Yes | MSMEG_5045 | MSMEG_3185 |
| C8_1 | 8 | 32 | 28 | Increase | High | Yes | MSMEG_4108,<br>MSMEG_4109, | MSMEG_1959,<br>MSMEG_1506,<br>MSMEG_3579,<br>MSMEG_4606,<br>upstream of<br>MSMEG_0520 |
| C8_2 | 8 | 32 | 28 | Increase | High | Yes | upstream of *nuoA* | MSMEG_1959,<br>MSMEG_3639,<br>MSMEG_5726,<br>MSMEG_6701,<br>MSMEG_3524,<br>MSMEG_6487,<br>upstream of<br>MSMEG_6223 and<br>MSMEG_5710 |
| C16_1, C16_2 | 16 | 64 | 56 | Increase | High | Yes | MSMEG_4064 | MSMEG_1959,<br>MSMEG_1957,<br>MSMEG_2254,<br>MSMEG_2174,<br>MSMEG_5283 |
| C16_L | 16 | 64 | 56 | No change | High | Yes | MSMEG_5045 | MSMEG_6133 |
| Increasing drug profile |  |  |  |  |  |  |  |  |
| R1 | 1-16 | 64 | 28 | Increase | High | Yes | No | MSMEG_5019,<br>upstream of<br>MSMEG_0573 and<br>MSMEG_0575 |
| R2 | 1-16 | 64 | 56 | Increase | High | Yes | No | MSMEG_1959,<br>MSMEG_5972, |
| R3 | 1-16 | 64 | 56 | Increase | High | Yes | No | upstream of<br>MSMEG_0573 and<br>MSMEG_0575 |
| WT_Ev | 1-16 | 64 | 28 | No change | High | Yes | No | MSMEG_6225,<br>MSMEG_5747 |

Among the diverse set of mutations identified, mutation in the gene *lfr*R was found to be conserved in all the evolved populations. This gene encodes for a TetR-family transcriptional regulator, LfrR which is involved in negative regulation of the efflux pump LfrA, known for extrusion of fluoroquinolone drugs along with other substrates such as Ethidium Bromide (EtBr), anthracyclines etc. thereby, promoting intrinsic antibiotic resistance in *M. smegmatis* (Figure 5.A) (38). The LfrR protein is a 189-residue long homodimer and comprises an N-terminal DNA binding domain (DBD) from residues 10-51 and a C-terminal ligand binding and dimerization domain (LBD). Although all evolved populations harbored mutations in the *lfr*R gene, in most of the populations, the mutation was an insertion resulting in a truncated protein with loss of the protein dimerization domain (a part of C-terminal ligand binding domain) while other domains being unaffected. Populations C8_1, C8_2 and C2_L harbored an SNP in the N-terminal DNA binding domain while C4_1 and C16_L strain had an SNP in the protein dimerization domain. The C2_2 population harbored an INDEL in the NTD leading to complete loss of protein. The C2_L population harbored an SNP just outside of the DBD of the LfrR protein while the WT_Ev population exhibited the same *lfr*R mutation as in the C16 population (insertion in the protein dimerization domain) (Figure 5.B).

**Figure 5.**
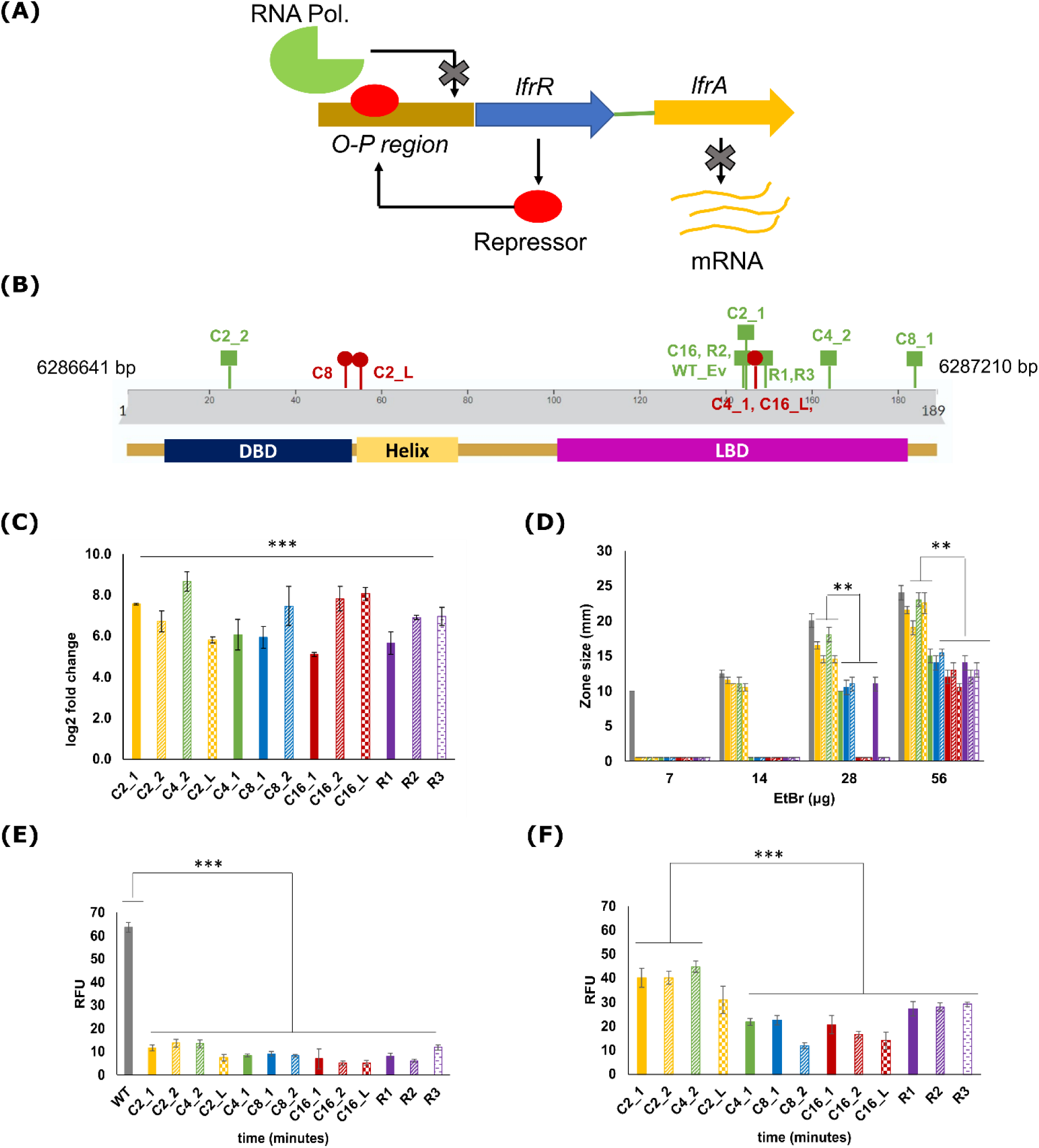
*lfr*A mediated resistance in the evolved populations. (A) Schematic for regulation of *lfr*A gene expression by the *lfr*R gene. (B) Schematic representation of LfrR protein sequence with different domains and the position of different mutations in the evolved populations. Green square markers represent Insertions while red round markers represent SNPs. Bottom bars represent different protein domains and their position in the protein: blue bar represents the N-terminal DBD, yellow bar represents the external helix to which DBD is attached and the pink bar represents C-terminal LBD. (C) *lfr*A gene expression level in different evolved populations with respect to the WT strain. 16S has been used as the housekeeping gene. (D) EtBr susceptibility testing using disc diffusion assay for the different evolved populations (Limit of detection: 10 mm). End point intracellular EtBr levels in the evolved populations at (E) 3.5 µg/ml and (F) 14 µg/ml EtBr concentration after 1 hour incubation. Standard deviation has been calculated over triplicate samples. * Indicates P < 0.05, ** indicates P < 0.01 and *** indicates P < 0.001. The C2 colonies are in yellow, the C4 is green, C8 in blue and the C16 in red. Purple represents the Ramp colonies.

All the different *lfr*R mutations led to over-expression of the LfrA efflux pump in the evolved populations (Figure 5.C). Consistent with efflux pump over-expression, all evolved colonies also exhibit increased resistance to EtBr (Figure 5.D). To understand the difference in sensitivity across the evolved populations, intracellular EtBr levels were measured using a fluorescence-based assay (details in methods). EtBr being a fluorophore and a common substrate for most of the efflux pumps, was an ideal candidate for measuring intracellular accumulation levels in the evolved populations. At a sub-inhibitory concentration of 3.5 µg/ml, all the evolved populations exhibited negligible EtBr accumulation (Figure 5.E). However, at higher concentrations of 14 µg/ml, the more sensitive populations (C2_1, C2_2 and C4_2) accumulated higher EtBr with respect to all the other populations (Figure 5.F). Thus, the increase in resistance of the evolved colonies can be attributed to the increased expression of *lfr*A gene. However, although EtBr MIC was clearly correlated to intracellular EtBr levels, no correlation was seen between MIC and efflux pump gene expression levels in the low and high mutants (Supplementary file S2, figure S1.A). To determine if the increased MIC of Norfloxacin was also due to reduced intracellular levels, intracellular level of Norfloxacin was also measured for few representative populations (Supplementary file S2, figure S1.B), where levels were found to be lower compared to WT.

### *lfr*R over-expression restores susceptibility in the evolved mutants

All the evolved populations harbored a mutation in the *lfr*R gene, leading to its inactivation as suggested by the over-expression of *lfr*A gene. Thus, to estimate the contribution of the different *lfr*R mutations on the resistance acquired, we expressed additional functional copies of the *lfr*R gene in WT strain and the different evolved populations. The WT *M. smegmatis* strain with extra *lfr*R copies was named as WT-OE. Similarly, *lfr*R over-expression was also carried out in one of the representative colonies from the different evolved populations. *lfr*R over-expression led to a 2-fold increase in Norfloxacin sensitivity in case of the WT-OE strain. Norfloxacin sensitivity also increased in all the other OE strains with levels similar to WT levels upon over-expression of *lfr*R (Figure 6.A and 6.B).

**Figure 6.**
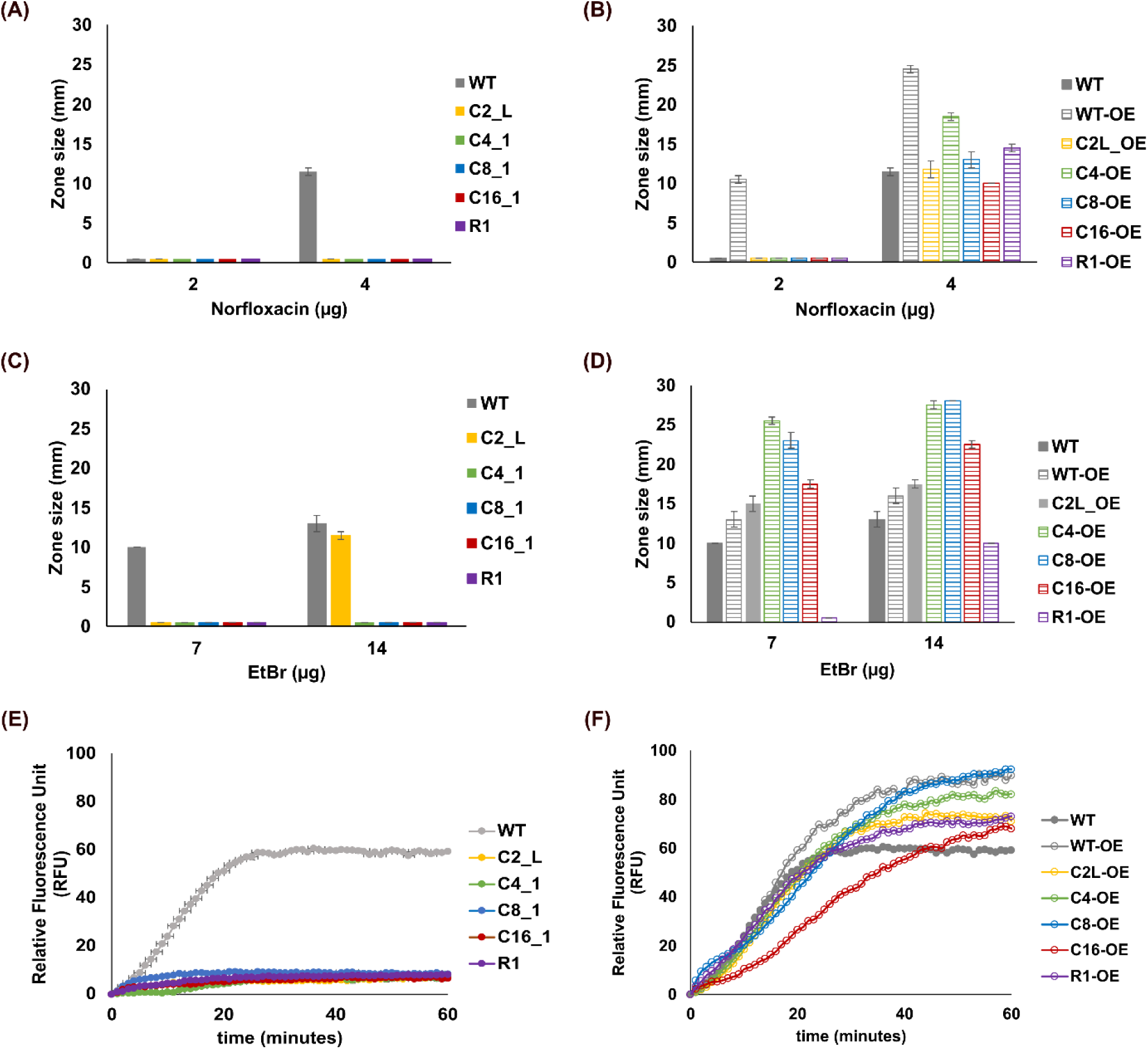
Phenotypic characterization of the evolved mutants over-expressing *lfr*R gene. (A) and (C) represent susceptibility testing for Norfloxacin and EtBr respectively against the representative evolved colonies while (B) and (D) represent susceptibility testing for Norfloxacin and EtBr respectively against the representative colonies with *lfr*R over-expression (lfrR-OE strains) (Limit of detection: 10 mm). (E) and (F) represent intracellular EtBr accumulation for the parent colonies and the lfrR-OE colonies at EtBr concentration of 3.5 µg/ml. Error bars have been calculated over triplicate samples. * Indicates P < 0.05, ** indicates P < 0.01 and *** indicates P < 0.001.

WT-OE strain also exhibited a two-fold increase in sensitivity to EtBr. Further, EtBr sensitivity was restored to WT levels in all the mutants except R1-OE where a 2-fold increase in sensitivity was observed (Figure 6.C and 6.D). In accordance with the increased sensitivity, all the lfrR-OE strains exhibited significant increase in EtBr accumulation as compared to their parent populations (Figure 6.E and 6.F).

### *lfr*R mutations are the first mutation to appear but not enough to impart high-level resistance

Given the conservation of *lfr*R mutations in populations evolved under diverse evolutionary profiles, we hypothesized that these were the first mutations to appear and get fixed in the population at high frequency. We therefore sequenced the full length *lfr*R gene from the initial passages of a few representative evolved lineages including C4_1, C4_2, C16_1 and C2_L population. The *lfr*R mutation was present in all the samples sequenced, confirming our hypothesis. In C4_1 and C4_2 colonies, the mutation identified in P1 was the same as that observed in the end point isolate. However, the C2_L population and the C16_1 colony harbored a different mutation in the initial passage. Details of *lfr*R mutations identified in different passages in the representative lineages is provided in Supplementary file S2, Table S3.

Being the most conserved mutation and the first mutation to appear during the evolutionary trajectory, it became imperative to quantify its role in imparting resistance. In a previous work from our lab, it was shown that *lfr*A over-expression can lead to a maximum EtBr MIC of 14 µg/ml (Barnabas et al. 2022). However, in many of our high resistance mutants, EtBr MIC of 28-56 µg/ml has been observed indicating potential role of other resistance mechanisms. To understand the factors behind the increased EtBr MIC of greater than 14 µg/ml, the evolved mutant colonies, C4_1 and C4_2 were chosen as they differed significantly in their resistance and intracellular accumulation levels despite over-expression of *lfr*A in both. Also, C4_1 colony had just one additional significant mutation in MSMEG_5045 (other two mutations were in pseudogenes) apart from the *lfr*R mutation. Since the same *lfr*R mutation was present in passage 1 and remained constant till passage 20, we tested intermediate passage strains for estimation of MIC and intracellular EtBr levels.

Passage 1 of C4_1 strain (C4_1_P1) displayed lower MIC compared to the subsequent passages for both Norfloxacin and EtBr and was the same as the C4_2 strain, passage 20 (C4_2_P20). In accordance with the MIC values, intracellular accumulation levels were significantly higher in C4_1_P1 compared to the other intermediate passage strains. Remarkably, the increased accumulation observed in Passage 1 strain was also similar to C4_2_P20 strain. This clearly indicated that the C4 population had an initial low resistance which gradually increased owing to the decreased accumulation. However, this high-level resistance obtained from C4_1 passage 5 (C4_1_P5) could not be attributed to *lfr*R mutation alone. WGS analysis of C4_1_P1 and C4_1_P5 strain revealed the presence of mutation in MSMEG_5045 only in C4_1_P5 strain which correlated well with the increased MIC and decreased accumulation in that strain (list of unique mutations in C4_1_P1 strain and C4_1_P5 strain is provided in supplementary file S1). A schematic of resistance profile and mutations identified in intermediate passages of C4_1 colony is provided in Figure 7.

**Figure 7.**
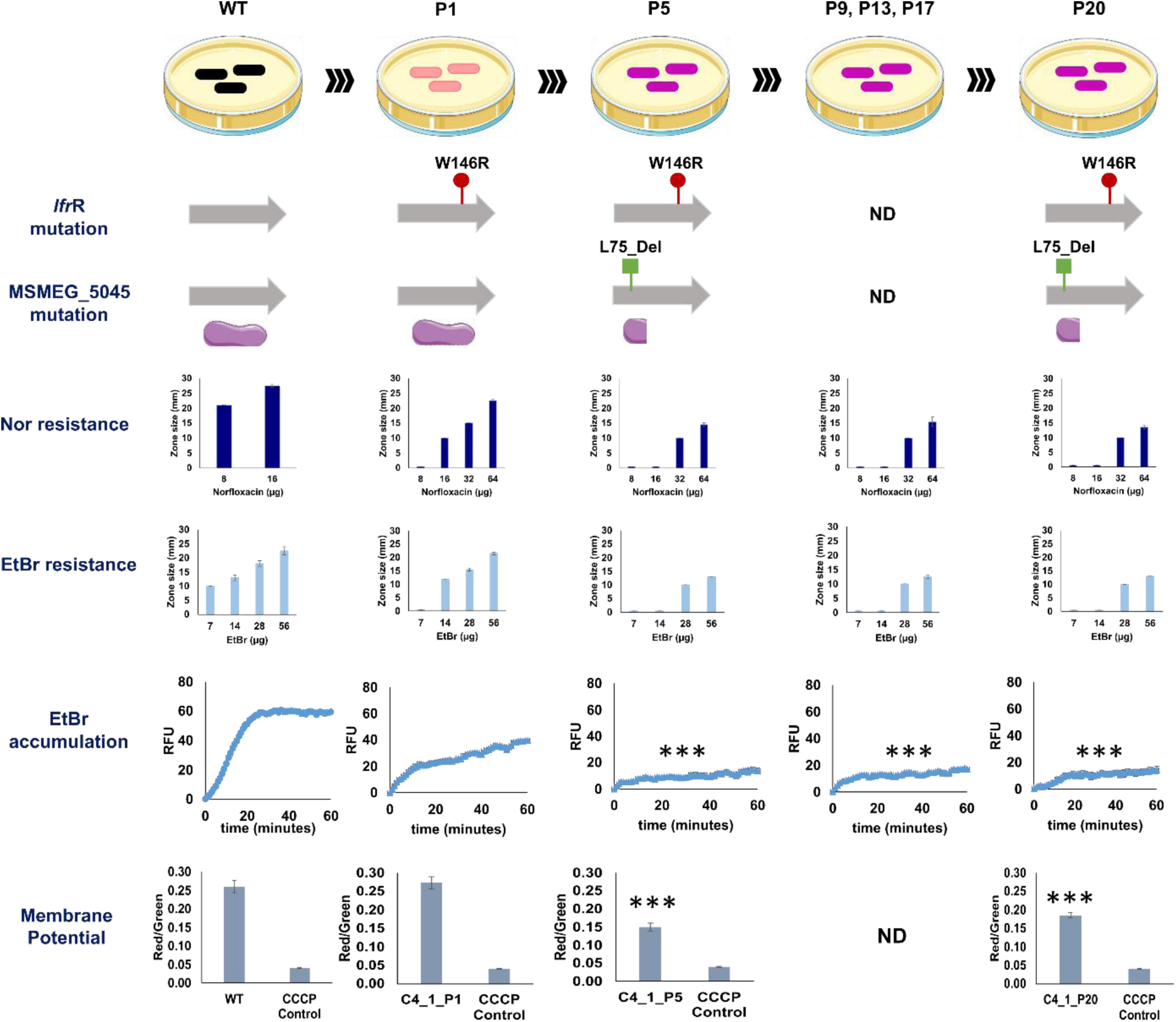
Sequential accumulation of mutations in C4_1 strain and its impact on resistance. Comparison of C4_1 intermediate passage strain (C4_1_P1, C4_1_P5, C4_1_P9, C4_1_P13, C4_1_P17) with the end point strain (C4_1_P20) in terms of the mutations identified, drug susceptibility and intracellular EtBr accumulation. Same *lfr*R mutation is conserved in all the C4_1 passage from P1 to P20 (red bars on top of the gene indicate SNP). Mutation in MSMEG_5045 appeared in P5 strain and the same mutation was conserved in P20 strain (green bars on top of the gene indicate deletion leading to formation of truncated protein represented in purple). Increase in resistance for both Norfloxacin and EtBr was observed from P5 onwards and sustained till P20. The increase in EtBr resistance also correlated with the decreased intracellular EtBr accumulation and decreased membrane potential from P5 onwards. Error bars have been calculated over triplicate samples. * Indicates P < 0.05, ** indicates P < 0.01 and *** indicates P < 0.001. Significance of p-value has been calculated with respect to the WT strain.

Acquisition of a mutation in MSMEG_5045 (coding for a dehydrogenase enzyme) led to increased resistance. There are previous studies reporting the role of dehydrogenase gene mutations in increasing resistance (39–41). Since, the dehydrogenase genes are known to impact pathways requiring energy translocation and hence, are likely to affect the efflux activity and membrane potential in cells, we further determined membrane potential of various C4_1 strains While C4_1_P1 had membrane potential similar to that of WT, the membrane was depolarized in C4_1_P5 and C4_1_P20 strains, coinciding with the appearance of the mutation in MSMEG_5045 gene encoding for a dehydrogenase. Since the mutation in MSMEG_5045 led to the formation of a truncated protein, we carried out phenotypic characterization of the Δ5045 strain. Absence of MSMEG_5045 led to a four-fold increase in EtBr MIC, decreased EtBr accumulation and reduced membrane potential (Supplementary file S2, Figure S2.A, S2.B and S2.C). Interestingly, the decrease in accumulation observed in Δ5045 strain was somewhere between the WT strain and the evolved mutants. Thus, we hypothesize that mutation in MSMEG_5045 leads to high-level resistance in combination with the pre-existing *lfr*R mutation.

### Selection of *lfr*R mutations is largely background dependent

Mutation in the *lfr*R gene was conserved across all the evolved populations irrespective of the evolution profile. To confirm that these mutations are selected primarily to relieve the repression of the LfrA pump, we evolved a *lfr*A deletion strain (Δ*lfr*A) under Norfloxacin selection pressure as three independent lineages (details in methods) and the evolved population was named as Δ*lfr*A_Ev. The WT_Ev population has been used as the control population here. Post evolution, the evolved mutants were characterized phenotypically for growth, fitness, and sensitivity to both Norfloxacin and EtBr. The Δ*lfr*A_Ev population exhibited increased resistance to both Norfloxacin and EtBr with a 16X and 32X increase in MIC respectively. To identify resistance determining factors, the QRDR regions of *gyr*A and *gyr*B along with the full length *lfr*R gene were sequenced from all three lineages of Δ*lfr*A_Ev population. None of the three replicate populations had any mutation in either of the three genes sequenced. Thus, these results confirm that the selection of *lfr*R mutations is strongly dependent on the genetic background of the evolved strains (presence of *lfr*A gene). Data for evolution profile and phenotypic characterization of Δ*lfr*A_Ev strain is provided in Supplementary file S2, figure S3.

## Discussion

Despite the recent advancements in the field of science and technology, antimicrobial resistance evolution in bacteria has been a global threat claiming millions of lives worldwide. Although many ALE experiments have been carried out to decipher the resistance determinants, most of these experiments take into consideration resistance emergence under a given evolutionary profile. In this study, we investigate the correlation between drug profile and the resistance mechanism by taking into account different parameters affecting ALE such as drug concentration, mode of drug administration and the number of generations. Since, ALE outcome is dependent upon multiple drug and host parameters, combining different parameters would help to identify conserved resistance determinants and resistance mechanisms that could serve as anti-evolution targets to impede resistance evolution.

WT *M. smegmatis* strain was used as the model organism and exposed to a range of Norfloxacin concentrations from 0.25X MIC to 4X MIC. Parallel evolution experiments were carried out under different constant drug pressures ranging from sub-MIC to 4X MIC along with evolution under stepwise increasing drug pressure for varying numbers of generations. Evolution under diverse evolutionary profiles for up-to 4X Norfloxacin MIC gave rise to mutants with varying resistance levels without any drug target mutation. In the different Constant populations, the resistance level and the number of mutations increased proportionally to the selection pressure. Since the different Constant populations were evolved for the same number of generations, the increase in the number of mutations in C8 and C16 populations indicated an increased mutation rate as a function of the drug exposure. The mutation rates across the different populations ranged from 4.34*10^-10^ to5.64*10^-9^. In a previous incidence in clinical isolates of *E. coli,* increased mutation rate leading to development of high-level fluoroquinolone resistance has been observed (42). In another study on *E. coli* evolved under Norfloxacin, a linear relationship was found between antibiotic concentration and the mutation rate (43). Although no mutations in specific mutator genes were identified in any of the evolved strains in this study, the increased mutation rates could possibly be due to alteration in expression of genes involved in DNA damage response pathway or other general stress response pathway as seen previously (44).

Increase in resistance due to sequential accumulation of mutations has been reported previously under laboratory evolution experiments (45–48) as well as within host (49, 50). We also observed sequential addition of mutations in the different evolved populations. Mutation in a regulatory gene, *lfr*R, was conserved in all the evolved populations, irrespective of the evolutionary profile, Targeted sequencing of the *lfr*R gene further confirmed that these mutations are the first to appear under diverse evolutionary profiles. Emergence of efflux related mutations as the first defense mechanism in mycobacteria has been seen previously for Bedaquiline and Azithromycin (46, 51). However, divergence was observed in the set of genes being added after the initial *lfr*R mutation in the populations evolved under constant vs increasing drug profiles. While repeated exposure of the same drug concentration led to enrichment of different dehydrogenase genes, successively increasing drug concentration selected for global regulatory genes. The fixation of regulatory genes under a fluctuating drug environment could be indicative of a more general stress response by the bacteria. Apart from the regulatory gene, *lfr*R, Ramp populations harbored mutations in and around other regulatory genes like MSMEG_5019 and MSMEG_0573. The gene MSMEG_0573 encodes an ECF sigma factor, RpoE1 which is known to regulate transcription under various stress responses (52). This indicated that efflux is selected as the primary resistance mechanism followed by divergent mechanisms facilitating the efflux-based mechanism.

The selection of *lfr*R mutations irrespective of the drug concentration exposed during evolution suggested these mutations to be more beneficial as compared to the drug target mutations under the evolutionary conditions in this work. The role of efflux pumps in strains without drug target mutations has been highlighted previously (25). The most common type of mutation was a nucleotide insertion leading to formation of a truncated protein while few other colonies exhibited SNPs in the *lfr*R gene. The resulting inactivation of the LfrR repressor led to over-expression of the *lfr*A gene across the different evolved mutants. The different *lfr*R mutations leading to efflux pump over-expression were in accordance with the selection bias for inactivating mutations selected in *mar*R regulator in Ciprofloxacin resistant *E. coli* clinical isolates (53). Efflux regulatory genes having SNPs have also been reported leading to efflux pump over-expression (54).

The dominant role of the mutations in *lfr*R gene was further confirmed when <u>o</u>ver-expression of the functional WT *lfr*R gene copy restored both Norfloxacin and EtBr sensitivity to WT levels in the different evolved populations. It is however important to take into consideration the fact that the background for *lfr*R over-expression was not completely devoid of the *lfr*R gene. Thus, to gain complete idea of the effect of different *lfr*R mutations, the *lfr*R gene copy with the various mutations should be expressed in a *lfr*R deletion background.

The data shows that mutations in *lfr*R alone can only lead to a 2-fold increase in MIC indicating the role of other mutations added later. Repeated exposure of the same drug pressure, as seen in the constant populations, allowed selection of a mutation in a dehydrogenase gene (MSMEG_5045) in C4_1 strain as early as passage 5 indicating it to be the next mutation after *lfr*R. Other constant populations displaying high resistance phenotypes also harbored mutations in different dehydrogenase genes (Table 1). The C8_1 population harbored many mutations in *pnt*A/B genes encoding transhydrogenase. These enzymes play an important role in regulating the NADP^+^: NADPH redox homeostasis in both prokaryotes and eukaryotes (55, 56). These genes are also known to be involved in nicotinate and nicotinamide metabolism (https://www.kegg.jp/pathway/map=map00760&keyword=pntA). Nicotinamide adenine dinucleotide (NAD) is a coenzyme for numerous vital redox reactions in the cell such as glycolysis, TCA cycle and DNA synthesis to name a few (57). C8_2 populations on the other hand harbored a mutation in the *nuo*A gene (NADH dehydrogenase I). A study in *P. Aeruginosa* has reported downregulation of few of the *nuo* genes to be linked to increased resistance to certain antibiotics (58). A double deletion *Mycobacterium tuberculosis* mutant (Δ*ndh*Δ*nuo*AN) was found to be the most severely attenuated strain *in vivo* suggesting vital role of *nuo* genes in the absence of primary NADH dehydrogenases (*ndh*) (59). C16 population harbored multiple mutations in two genes encoding alcohol dehydrogenase, MSMEG_4032 and MSMEG_4046, both being reported to be upregulated under oxidative stress in *M. smegmatis* (60). The role of several dehydrogenase genes leading to increased isoniazid resistance in *M. tuberculosis* strain has been suggested previously (61). Since multiple dehydrogenase gene mutations were identified across different evolved populations, deciphering the role of all the individual gene mutations and the exact mechanism associated with resistance requires further investigation. However, we have shown that mutation in one of the dehydrogenase genes, MSMEG_5045, caused membrane depolarization which coupled with enhanced drug efflux led to high-level resistance acquisition. The reduced membrane potential could either exert its effect by reducing drug uptake, as has been reported previously for aminoglycosides (62), or may influence efflux pump activity A recent study has reported changes in membrane potential being linked to efflux pumps affecting bacterial physiology (63).

To conclude, this study highlights how drug profiles strongly influence the adaptive pathways adopted by bacteria to gain resistance. While efflux-based resistance through mutation in an efflux pump regulator was the first defense mechanism, subsequent mutations that were acquired depended on the evolutionary profile. The data also strongly suggested the role of dehydrogenase genes in augmenting the initial resistance, and thus could serve as targets for development of anti-evolutionary agents. Acquiring mutations in the genes encoding for drug targets, usually renders the drug ineffective. We show that mutations in the drug target genes are acquired only beyond a threshold concentration. While similar studies are required for other drugs, this knowledge can help design effective treatment protocols that prevent or delay the appearance of resistance.

## Materials and methods

### Bacterial strain and culture condition

*Mycobacterium smegmatis* mc^2^155 strain has been used as the Wild Type strain throughout the experiments and *E. coli* DH5α strain has been used for cloning experiments. Δ*lfr*A strain was received from Prof. Miguel Viveiros, Global Health and Tropical Medicine, GHTM, Instituto de Higiene e Medicina Tropical, IHMT, Universidade Nova de Lisboa, UNL, Lisboa, Portugal as a kind gift. Middlebrook 7H9 supplemented with 0.15% (v/v) Tween 80 and 0.44% (v/v) 100% glycerol has been used as the standard growth medium for *Mycobacterium smegmatis* mc^2^155 liquid cultures and LB broth for *E. coli* DH5α strain. For solid cultures, Luria-Bertani (LB) supplemented with 1.5% agar has been used as the growth medium for both the strains. All the culture media were purchased from Hi-Media, India. Continuous shaking at 200 rpm and 37°C was used as the ideal condition for bacterial growth. Antibiotics Norfloxacin and Ethidium Bromide (EtBr), were purchased from Sigma and Hi-Media, India respectively.

### Evolution of resistance mutants

. For generation of Constant and Ramp populations, Wild Type *M. smegmatis* strain (mc^2^155) was evolved on a range of constant Norfloxacin concentrations from 0.5X to 4X WT MIC and stepwise increasing Norfloxacin concentrations from 0.25X to 4X WT MIC for 330 generations respectively. Single isolated colony from the WT strain was streaked onto Luria-Bertani (LB) plates containing desired antibiotic concentration and incubated till independent colonies appeared on the plates before streaking onto the next plate with the same or higher antibiotic concentration till the 20th passage. Populations were evolved in duplicates for each of the evolution profiles along with a Control population evolved simultaneously without any drug pressure on plain LB plates.

Evolution in liquid media under Constant Norfloxacin concentration of 2 μg/ml and 16 μg/ml (in duplicates) was also done in a similar fashion for 20 passages with higher transfer size of 10^5^ cells at each passage. Following the same transfer size, evolution of WT_Ev and ΔlfrA_Ev strains was carried out ranging from their respective sub-MIC to 4X Norfloxacin MIC. Following the final passages, six colonies from each Constant and Ramp population were picked, re-suspended in M7H9 broth and 50% glycerol (v/v) and stored as glycerol stock for further use. For liquid evolved mutants, mid-log phase cells at each passage were stored as glycerol stock. Detailed protocol is provided in Supplementary file S3. List of the different evolved strains with description is provided in Supplementary file S2, Table S4.

### Measurement of growth kinetics

For measuring growth kinetics, 400 µL of the inoculum was taken from working stock and inoculated into 50 mL of M7H9 broth and incubated at 37° C and at 200 rpm in a shaker for ∼50 hours. Absorbance at 600 nm (OD_600nm_) measurements were taken at regular intervals which corresponded to bacterial growth. Absorbance values were plotted as a function of time from which doubling time of different bacterial populations was calculated such that **doubling time** equals to **ln (2)/slope of the graph**.

### Susceptibility testing

Susceptibility testing was done using the Zone of Inhibition (ZOI) test for the different evolved colonies. For this, 25 µL of cells (OD_600nm_ = 0.5) grown in M7H9 broth were streaked onto Luria Bertani (LB) agar plates. After this, wells were cut in the plates, a desired amount of antibiotic added to each well and plates incubated for 48 hours. Post incubation, zone size was measured and results recorded. Lesser zone size corresponded to increased resistance.

### Determination of Minimum Inhibitory Concentration (MIC)

MIC determination was done through broth microdilution assay using 96 well transparent sterile microplates (Thermo Fisher Scientific, USA) in which 200 µL of 10^6^ cells/mL (corresponding to OD_600nm_ = 0.05) were added in each well of the microplate. To each well, the desired antibiotic concentration was added followed by incubation at 37°C for 48 hours. Post incubation, five drops of 10 µl each from all the wells were dropped on LB agar plates which were further incubated for 48 hours at 37°C (Drop Count Method) and colony count done using the formula below:

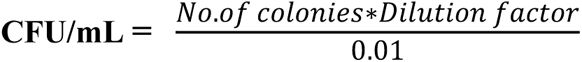

The minimum drug concentration which gave a CFU count of 10^6^ cells/ml or below was the Minimum Inhibitory Concentration (MIC).

### Intracellular accumulation and efflux studies using EtBr as tracer molecule

For EtBr uptake study, kinetic measurement of intracellular EtBr accumulation using a fluorescence-based method was used as mentioned in a previous paper (27). For efflux assay, cells were grown till mid-log phase and EtBr at the desired concentration with CCCP (10 µg/ml) was added to 1 ml of the cell suspension followed by 1 hour incubation at 37°C. The samples were then centrifuged at 10,000 RPM for 10 minutes at 4°C and washed and resuspended in 1 ml ice cold PBS as positive control and PBS with CCCP as negative control. 200 μL aliquot from each sample was then transferred to the 96 well plates followed by fluorescence measurement similar to accumulation assay (64).

### PCR amplification and Sanger sequencing

PCR amplification for the target genes, *lfr*R, *gyr*A and *gyr*B was done by using a standardized protocol using 500 ng of genomic DNA, gene specific forward and reverse primers (0.5 µM final concentration) and 0.025 % DMSO were added along with DreamTaq DNA polymerase (Thermo scientific), reaction buffer and dNTPs. The PCR reaction was performed in a thermal cycler (Hi Media) using optimized thermal cycling conditions. Sequencing of the amplified product was done using an automated DNA sequencer (Applied Biosystems 3730 DNA analyzer) by Eurofins Genomics India Pvt Ltd. Detailed protocol is provided in Supplementary file S3. A list of primers used for targeted sequencing of different genes is provided in Supplementary file S2, Table S5.

### Estimation of Membrane potential

Changes in the potential across the cell membrane were quantified by the 3,3′-Diethyloxacarbocyanine iodide (DiOC_2_, Sigma-Aldrich) dye (65). 0.5 OD cells were centrifuged and resuspended in PBS (without K^+^ ions) with 30 µM DiOC_2_. Samples were further incubated in dark for 1 hour at 37°C with aeration at 200 rpm. Post incubation, an aliquot of 200 µL of the sample was added in the 96 well plate and the fluorescence detected by excitation at 488 nm and emission at 520 nm and 620 nm for green and red fluorescence respectively. Data was plotted as the ratio of red/green fluorescence for each sample (66).

### Whole Genome Sequencing (WGS) studies

Evolved populations from their glycerol stocks were re-inoculated into M7H9 media, grown till mid log phase and genomic DNA isolated using a method adapted from Wright *et. al*. with minor modifications (67). The isolated DNA was sent out for sequencing to BGI Sequencing Services, Hong Kong, China and Novogene (HK) Company Limited. Sequencing was done on Illumina Hi-Seq X Ten, PE151 (average read length of 300 bp, average base coverage depth of 181, mapping efficiency of 99.9% and a total of ∼10 million reads).

The raw data files were uploaded onto the Galaxy web platform and further analysis was done using the public server at usegalaxy.org (68). Raw reads were mapped to the reference genome of *Mycobacterium smegmatis* mc^2^155 (GenBank accession number NC_008596.1) using Bowtie2. Elimination of duplicate reads was done using Mark Duplicates tool.

For variant identification (both SNPs and INDELs), VarScan tool was used and the following parameters were set: minimum coverage depth of 8X, minimum variant frequency of ≥0.01, minimum average base quality of ≥20 and a Phred score of ≥Q20 (>99% accuracy) (69, 70). Evolved strains are compared with control strain and mutations having frequency of ≥ 0.3 and coverage of ≥ 50 were selected for further study.

### Measurement of gene expression

Gene expression studies have been done using CFX96 Touch^TM^ Real Time PCR Detection System (Bio-Rad, USA) for *lfr*A gene, in WT and the evolved colonies. RNA isolation was done using a Tri-reagent (Merck) based protocol (71) followed by cDNA synthesis using random primers and Reverse Transcriptase (RT) enzyme (ThermoFisher Scientific). Using 100 ng of cDNA as the template and 0.5 µM of sequence specific primers, qRT-PCR reaction was set based on a set protocol to identify differential expression levels of the genes in WT and the mutants. Detailed protocol is provided in Supplementary file S3.

The 2^-ΔΔct^ method (72) was used for differential gene expression analysis where 16S rRNA was used as the housekeeping gene:

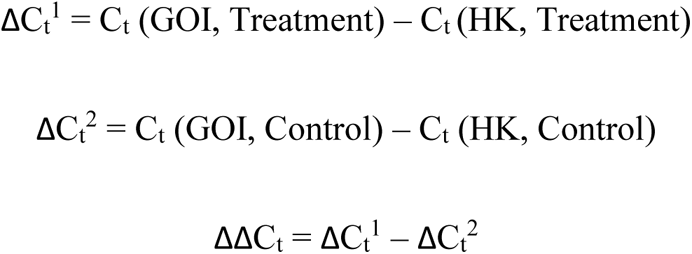

Relative fold change gene expression = **2^-ΔΔC**_t_ where, GOI is gene of interest, HK is house-keeping gene and C_t_ is threshold cycle. List of primers used for expression studies is provided in Supplementary file S2, Table S5.

### Generation of *lfr*R over-expression strains

For generation of overexpression strains, the *lfr*R gene was amplified using PCR from WT *M. smegmatis* mc2155 (NC_008596.1) genomic DNA. The amplified PCR product was sequentially digested using enzymes BamHI and XbaI and ligated into a multicopy plasmid pST-K, digested previously using the same restriction enzymes. pST-K is a replicative plasmid in *E. coli* DH5α strain used for constitutive expression of the gene inserted by virtue of P-myc-tetO promoter. Following generation of successful clones in *E. coli,* plasmid containing *lfr*R gene was transformed into *M. smegmatis* cells (WT strain and different evolved strains) via electroporation and selection of successful clones was done on LB plates supplemented with kanamycin (25 µg/ml). Nomenclature of the clones was done with OE as a suffix for the parent strain (WT-OE, C4-OE and so on). List of primers used for generation of OE strain is provided in Supplementary file S2, Table S5.

### Statistical analysis

Statistical analysis was performed using at least three biological replicates. Significance was determined using Student’s *t* test taking equal variances, where * indicates *P* < 0.05, ** indicates *P* < 0.01 and *** indicates *P* < 0.001.

## Supplementary material

**Supplementary file S1**: List of all unique mutations identified in the evolved populations

**Supplementary file S2**: Additional tables and figures

**Supplementary file S3**: Detailed methods

## Acknowledgement

The work was funded by the Department of Science and Technology (DST), India (SERB No. EMR/2016/007667). Akanksha would like to acknowledge the Department of Biotechnology (DBT) India and Industrial Research and Consultancy Centre (IRCC), IIT-Bombay for providing the fellowship.

## Author contribution

Conceived and designed the experiments: Akanksha, SM. Performed the experiments: Akanksha. Analyzed the data: Akanksha, SM. Wrote the paper: Akanksha, SM.

## Data availability

The raw genome sequence for the 19 evolved strains has been deposited in the NCBI database (BioProject ID: PRJNA956056).

